# Volumetric Scattering Microscopy

**DOI:** 10.64898/2026.04.03.716429

**Authors:** Zijun Gao, Keyi Han, Zhi Ling, Hongmanlin Zhang, Edward A Botchwhey, Wen-hao Liu, Xuanwen Hua, Shuyi Nie, Shu Jia

## Abstract

Optical scattering in biological tissues fundamentally limits fluorescence imaging by disrupting spatial and angular information, thereby restricting volumetric visualization. Although hardware-intensive and computational approaches have advanced scattering microscopy, practical three-dimensional imaging through tissue remains constrained by instrumental complexity and axial ambiguity. Here, we present volumetric scattering microscopy (VSM), a scan-free, optical–computational framework for three-dimensional fluorescence imaging in scattering biological media. VSM captures angularly resolved speckle-encoded fluorescence using an aperture-segmented Fourier light-field configuration and reconstructs volumetric structure through adaptive feature-based descattering and joint sub-pupil alignment. This hybrid strategy preserves angular information embedded in scattered light without wavefront measurement or mechanical scanning, while maintaining the simplicity of a standard epi-fluorescence architecture. We demonstrate high-fidelity volumetric reconstruction across phantoms, engineered cellular systems, *ex vivo* tissues with volumetric muscle loss, and intact *Xenopus* embryos, achieving preserved spatial resolution, enhanced optical sectioning, and quantitative accuracy under strong scattering conditions. VSM supports large-field, robust volumetric imaging in both layered and fully embedded scattering environments. By transforming scattered light into a structured encoding resource, VSM establishes a scalable pathway toward routine three-dimensional fluorescence imaging in complex biological systems.

## INTRODUCTION

Fluorescence microscopy has fundamentally transformed modern biomedical science by enabling direct visualization of biological systems with high sensitivity, molecular specificity, and spatiotemporal precision^1,2^. As biological questions continue to expand in complexity, scale, and dimensionality, sustained advances in optical instrumentation and computational algorithms have progressively expanded imaging capabilities, reinforcing the indispensable role of fluorescence microscopy in biomedical research. From a physical standpoint, however, optical microscopy remains primarily constrained by two fundamental limits: diffraction, which bounds spatial resolution under ballistic light propagation, and optical scattering, which restricts imaging depth and information fidelity in heterogeneous biological media.

In fact, most fluorescence microscopy techniques rely on the ballistic propagation of emitted photons along well-defined optical paths to form high-fidelity images at the detector, a regime in which recent advances have enabled increasingly detailed observation of dynamic biological processes^3,4^ and nanoscale structures^5–8^. Biological tissues, in contrast, are intrinsically heterogeneous and exhibit spatially varying refractive indices, leading to strong, unpredictable photon scattering^9,10^. Consequently, ballistic photons are rapidly attenuated over depths of only a few mean free paths, typically on the order of hundreds of micrometers, resulting in severe degradation of image contrast and resolution when targets are embedded beneath scattering layers^11,12^. Overcoming optical scattering, therefore, remains a central yet incompletely resolved challenge for fluorescence microscopy, particularly in deep-tissue and *in situ* imaging contexts.

Over the past two decades, a wide range of strategies have been proposed to mitigate the limitations imposed by optical scattering in biological tissues^12–14^. For example, optical clearing techniques suppress tissue-induced scattering by chemically homogenizing refractive-index heterogeneity, thereby enabling deep, high-contrast optical microscopy^15–17^. Despite their effectiveness, the processing required for tissue clearing generally precludes live imaging and limits applicability to *in situ* studies. Beyond sample modification, considerable effort has been devoted to hardware-based imaging strategies that actively manipulate light propagation through scattering media. These advances, such as nonlinear excitation^14,18,19^, ballistic photon gating^20–22^, acoustic guiding^23–26^, wavefront shaping^27,28^, and transmission matrix measurement^29–31^, have demonstrated substantial gains in penetration depth, often exceeding an order of magnitude in the scattering mean free path. Moreover, these strategies have been extended to three-dimensional scattering imaging by leveraging time-gated detection and holographic phase measurements^32,33^. Nevertheless, the practical adoption of these techniques remains constrained by their reliance on complex optical architectures, high-cost components, and, in many cases, limited acquisition speed.

In parallel, computational approaches that require minimal sample preparation or hardware modification have attracted growing attention for their ease of implementation and adoption^34^. A prominent class of such strategies exploits the intrinsic robustness of speckle-encoded measurements to tissue-induced aberrations by leveraging the optical memory effect (OME)^35–37^. This phenomenon preserves short-range spatial correlations in scattered light, enabling object recovery via approaches such as speckle autocorrelation without requiring explicit knowledge of the scattering wavefront^38,39^. More recently, scattering compensation has been further advanced through a range of complementary strategies that extend imaging performance across sample density, computational efficiency, resolution, and field of view, including reflection-matrix methods^40,41^, learning-based approaches^42–45^, speckle illumination schemes^46–48^, and dimensionality-reduction frameworks^49–51^. However, scattering-induced depth insensitivity causes volumetric information to overlap within a single measurement, progressively degrading axial distinguishability. Consequently, despite their strengths, current computational approaches remain largely confined to two-dimensional imaging, and practical solutions for three-dimensional scattering microscopy remain underexplored^52–54^.

Here, we introduce volumetric scattering microscopy (VSM), an integrated optical and computational framework for scan-free, three-dimensional fluorescence imaging through scattering biological media. Specifically, VSM captures angularly resolved scattered fluorescence using an aperture-segmented Fourier light-field configuration, providing spatial-frequency recordings that support volumetric reconstruction. Each elemental image undergoes a unified processing pipeline that combines adaptive feature representation, collective sub-pupil alignment, and patch-wise volumetric deconvolution, thereby mitigating scattering-induced distortions and yielding a high-resolution 3D reconstruction. We demonstrate that VSM achieves high data efficiency and robust performance with minimal hardware complexity across diverse biological systems, including tissue specimens and whole organisms. Together, these results establish VSM as a practical and versatile approach for advancing fluorescence microscopy in complex scattering environments with broad relevance to biomedical research.

## RESULTS

### Principle and Framework of VSM

The principle underlying VSM is that, despite the apparent randomness of speckle-encoded scattering measurements, they preserve critical angular information that implicitly encodes volumetric structure and has largely remained unexploited^55^. Building on this insight, VSM departs from existing approaches by introducing a scan-free, physics-informed framework for three-dimensional fluorescence imaging through scattering media (**Supplementary Tables 1-2**). Experimentally, VSM employs speckle-encoded illumination and detection based on a customized upright Fourier light-field configuration, in which segmentation of the system pupil enables moderately aliased angular sampling of scattered fluorescence, thereby enhancing optical sectioning, while maintaining high photon efficiency with minimal alignment complexity^56,57^ (**Fig. 1a, Supplementary Fig. 1**).

**Figure 1.**
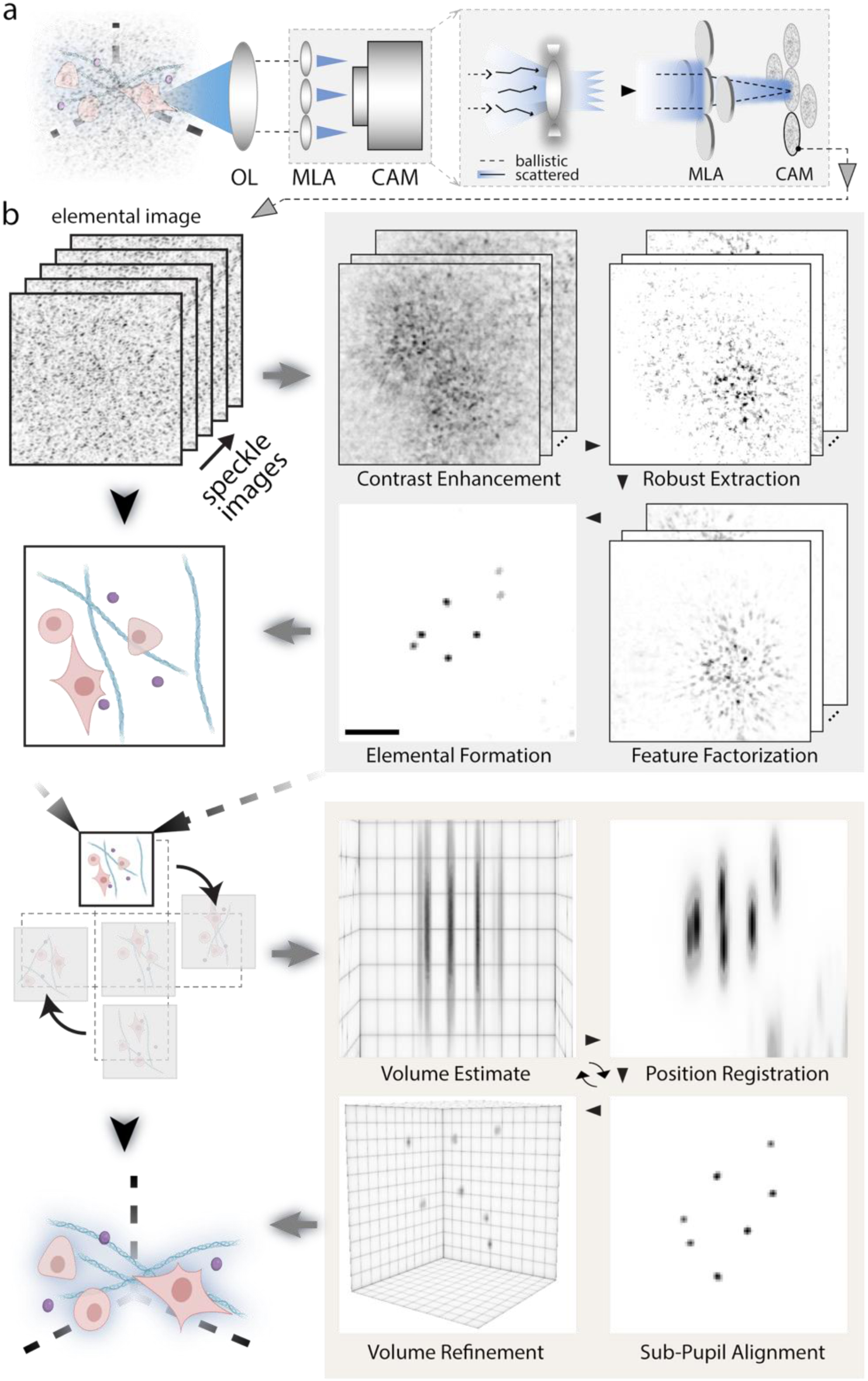
Volumetric scattering microscopy (VSM). **(a)** Schematic of the VSM setup, capturing angularly resolved scattered fluorescence using an aperture-segmented Fourier light-field configuration. The inset shows aliasing of spatial frequencies due to scattering, which facilitates enhanced optical sectioning. OL, objective lens; MLA, microlens array. **(b)** *Left panel*, schematic of the VSM algorithmic pipeline, including adaptive feature representation for each elemental image, sub-pupil alignment of all elemental images, and collective patch-wise volumetric deconvolution, for high-resolution 3D reconstruction. *Right panel*, detailed step-by-step illustration of processing modules (see details in **Supplementary Notes 2-4**), exemplifying scattering imaging of 2-μm fluorescent microspheres volumetrically distributed in agarose gel and beneath 3M tape using VSM. Scale bar: 100 μm.

Next, the acquired scattering-encoded light-field measurements are processed by VSM through a two-stage reconstruction pipeline (**Fig. 1b**). *First,* each elemental image undergoes adaptive, scattering-aware feature processing designed to accommodate distinct scattering regimes. In particular, under strong-scattering conditions, VSM provides a robust non-negative principal matrix factorization framework for addressing non-sparse signals and redundant background interference (**Methods**). In practice, each elemental image is preprocessed using Fourier-domain filtering to suppress high-frequency noise and background fluctuations^49^, then decomposed into sparse features and low-rank background components, followed by dimensionality reduction and feature assignment^58^ (**Fig. 1b, Supplementary Notes 2-3**). This feature-based processing exhibits strong robustness in highly scattering environments and thereby preserves angular information essential for volumetric reconstruction, outperforming conventional OME-based speckle autocorrelation and matrix-based approaches^58^. *Second*, the feature-represented elemental images are integrated into a three-dimensional volume through a collective sub-pupil alignment performed in parallel with patch-wise volumetric reconstruction (**Fig. 1b**). Because scattering disrupts the defined spatial relationships expected under ballistic propagation, descattering of individual elemental images does not preserve consistent positional correspondences across sub-pupils^50^. To address this challenge, VSM recovers the segmented-pupil geometry and conducts volumetric retrieval within a unified, concurrent, forward-backward iterative projection framework^59^, imposing positional constraints from a reference elemental image without requiring prior knowledge of sub-pupil misplacement (**Supplementary Note 4**). Collectively, this end-to-end pipeline enables efficient recovery and volumetric fusion of the elemental images and formation of high-resolution three-dimensional reconstructions, exhibiting rapid convergence, improved volumetric accuracy and resolution, and robust performance across imaging conditions (**Supplementary Figs. 2-5**).

### Characterization of VSM

We characterized VSM performance under representative scattering conditions using both phantom and biological samples. We first imaged 2-µm fluorescent microspheres distributed in an agarose gel, 1 mm beneath a layer of invisible tape (**Methods**), which produced scattering equivalent to approximately one scattering mean free path (1 ℓₛ)^58^. The raw elemental images exhibit fully developed speckle patterns with minimal discernible structural information (**Fig. 2a**). In contrast, VSM reconstructs high-quality elemental images that are consistent with the reference wide-field images acquired in the absence of the scattering layer (**Fig. 2b**) and subsequently recovers the three-dimensional distribution of the microspheres with high fidelity (**Fig. 2c**). Notably, the processing remains robust in the scattering environments, in comparison with conventional approaches (**Supplementary Fig. 6**). Quantitative analysis exhibited lateral and axial FWHM values of approximately 4 µm and 9 µm, respectively, in good agreement with the performance of the underlying Fourier light-field microscope (**Fig. 2d**).

**Figure 2.**
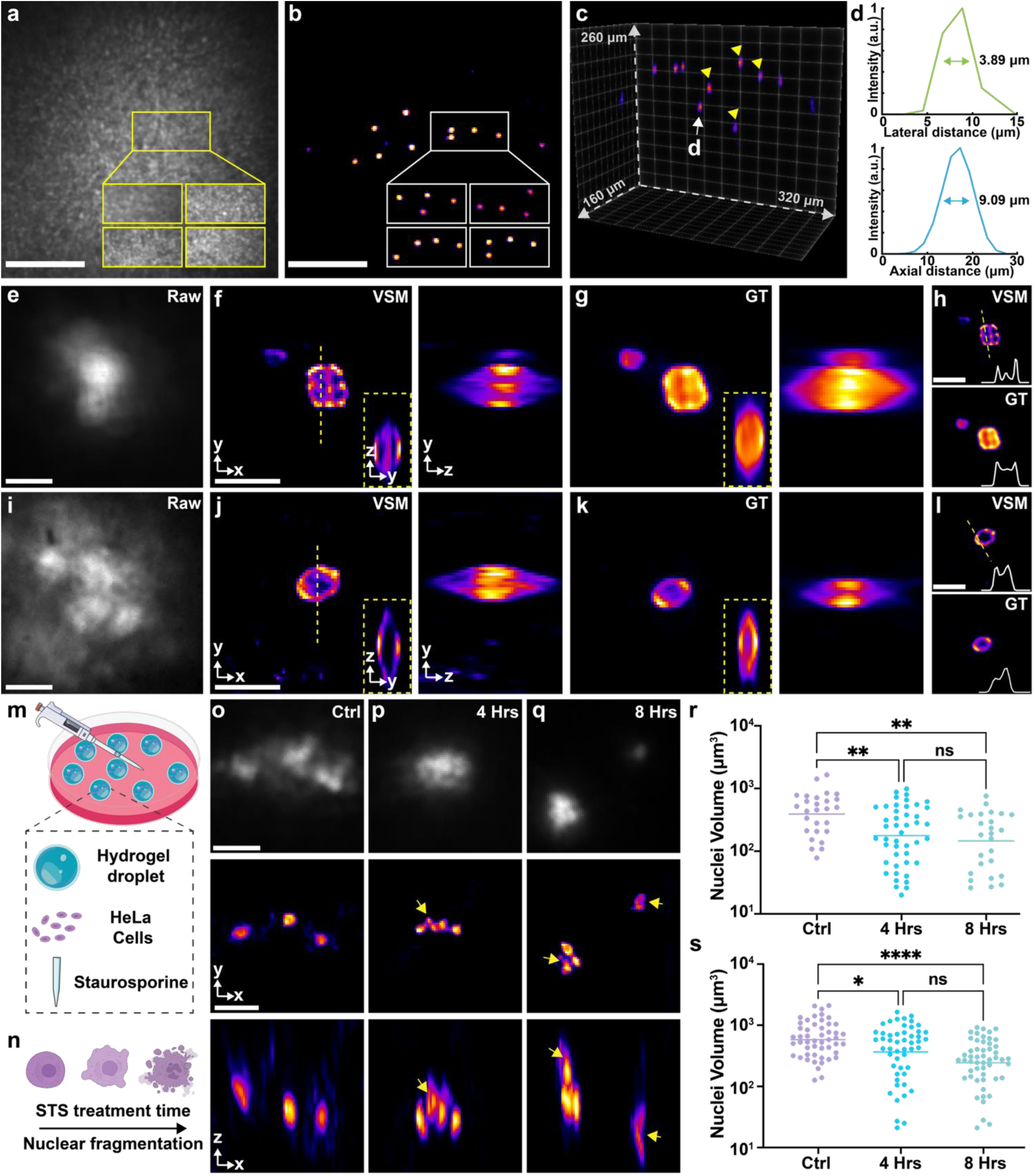
Characterization of VSM under phantom and biological scattering conditions. **(a, b)** Representative raw (a) and correspondingly descattered (b) elemental images of three-dimensionally distributed 2-µm fluorescent microspheres embedded in agarose and covered by invisible tape. Insets show the corresponding boxed regions of the object in other raw and descattered elemental images, where the speckle-encoded angular features are recovered. **(c)** Corresponding VSM reconstructed volume. Four yellow arrows indicate the boxed objects in (b). **(d)** 3D cross-sectional profiles of the microsphere, as marked in (c), exhibiting the lateral and axial FWHM values of 3.89 µm and 9.09 µm, respectively. **(e, f)** Representative raw elemental (e) and maximum-intensity projection (MIP) of VSM (f) images of a pollen grain placed beneath 150-µm-thick mouse skin. **(g)** Corresponding light-field reconstructed volume without the scattering layer. Insets in (f, g) show *x-z* cross-sections along the corresponding yellow dashed lines. **(h)** Cross-section images of the central layer, showing enhanced optical sectioning and resolution of fine details inside the pollen with VSM. **(i, j)** Representative raw elemental (i) and maximum-intensity projection (MIP) of VSM (f) images of three-dimensional hydrogel-cultured mesenchymal stem cells (MSCs) through 150-µm-thick mouse skin. **(k)** Corresponding light-field reconstructed volume without the scattering layer. Insets in (j, k) show *x-z* cross-sections along the corresponding yellow dashed lines. **(l)** Cross-section image of the central layer, showing enhanced optical sectioning of the cellular membrane with VSM. **(m, n)** Schematic of 3D-cultured HeLa cells embedded in alginate– Ca²⁺ hydrogel undergoing staurosporine (STS)-induced apoptosis and nuclear fragmentation. **(o-q)** Representative raw elemental (top) and VSM reconstructions in *x-y* (middle) and *x-z* (bottom) dimensions acquired through 150-µm-thick mouse skin at increasing STS treatment durations. Yellow arrows indicate progressive nuclear fragmentation resolved in both *x-y* and *x-z* views using VSM. **(r, s)** Quantitative nuclear volume distributions derived from VSM images (r) and conventional light-field results (s) over time, showing a consistent shift toward smaller fragmented nuclei during apoptosis progression. Statistical analyses were performed using the Kruskal-Wallis test followed by Dunn’s multiple comparisons test. ns, p > 0.05; *p ≤ 0.05; **p ≤ 0.01; ****p ≤ 0.0001. Scale bars: 100 µm (a, b), 50 µm (e-l, o-q).

We next extended VSM to samples with increased structural complexity and stronger scattering. Specifically, we imaged pollen grains placed 1 mm beneath a 150-µm-thick mouse skin section, which generated low-contrast speckle patterns due to both the dense internal structure of the pollen and the overlying scattering layer (**Fig. 2e**). As seen, VSM reconstructs the volumetric morphology (**Fig. 2f**) with strong correspondence to the ground truth (**Fig. 2g**). Notably, VSM exhibits enhanced optical sectioning compared to conventional light-field reconstruction (**Fig. 2h**), resulting from the improved sectional discrimination provided by aliased spatial frequencies owing to scattering that contributes to elemental image reconstruction (**Fig. 1a**). We further evaluated VSM using 3D hydrogel-cultured mesenchymal stem cells (MSCs) covered by the same skin layer. Despite the absence of discernible structure in the raw speckle measurements (**Fig. 2i**), VSM robustly reconstructs the hollow cell membrane morphology, preserving fine membrane curvature and lumen integrity (**Fig. 2j-l**), demonstrating the consistent performance in both reconstruction quality and optical sectioning.

Furthermore, we used VSM to image cell apoptosis in three-dimensional scattering microenvironments. HeLa cells were embedded in alginate-based hydrogel droplets (**Fig. 2m)** and captured through a 150-µm-thick mouse skin section, which recapitulates a physiologically relevant space (**Methods**). The 3D cell cultures were treated with staurosporine (STS), a protein kinase inhibitor, for varying durations to monitor STS-induced apoptosis progression via nuclear fragmentation (**Fig. 2n**). VSM overcomes strong tissue scattering, accurately recovering volumetric information from low-contrast speckle measurements and enabling clear resolution of these three-dimensional structural changes (**Fig. 3o-q)**. As observed, nuclei in the untreated control group remain intact and morphologically uniform (**Fig. 2o**), whereas after 4 and 8 hours of STS treatment, pronounced nuclear fragmentation is observed, characterized by multiple condensed nuclear fragments (**Fig. 2p-q**). Quantitative analysis of reconstructed nuclear volumes reveals a progressive decrease in average nuclear volume from 4,820 µm³ in controls to 2,146 µm³ after 8-hour treatment (**Fig. 2r**). Importantly, the volumetric trend measured by VSM closely matches the corresponding ground-truth measurements acquired without the scattering layer (**Fig. 2s**), demonstrating accurate recovery of apoptosis-associated morphological changes under scattering conditions.

**Figure 3.**
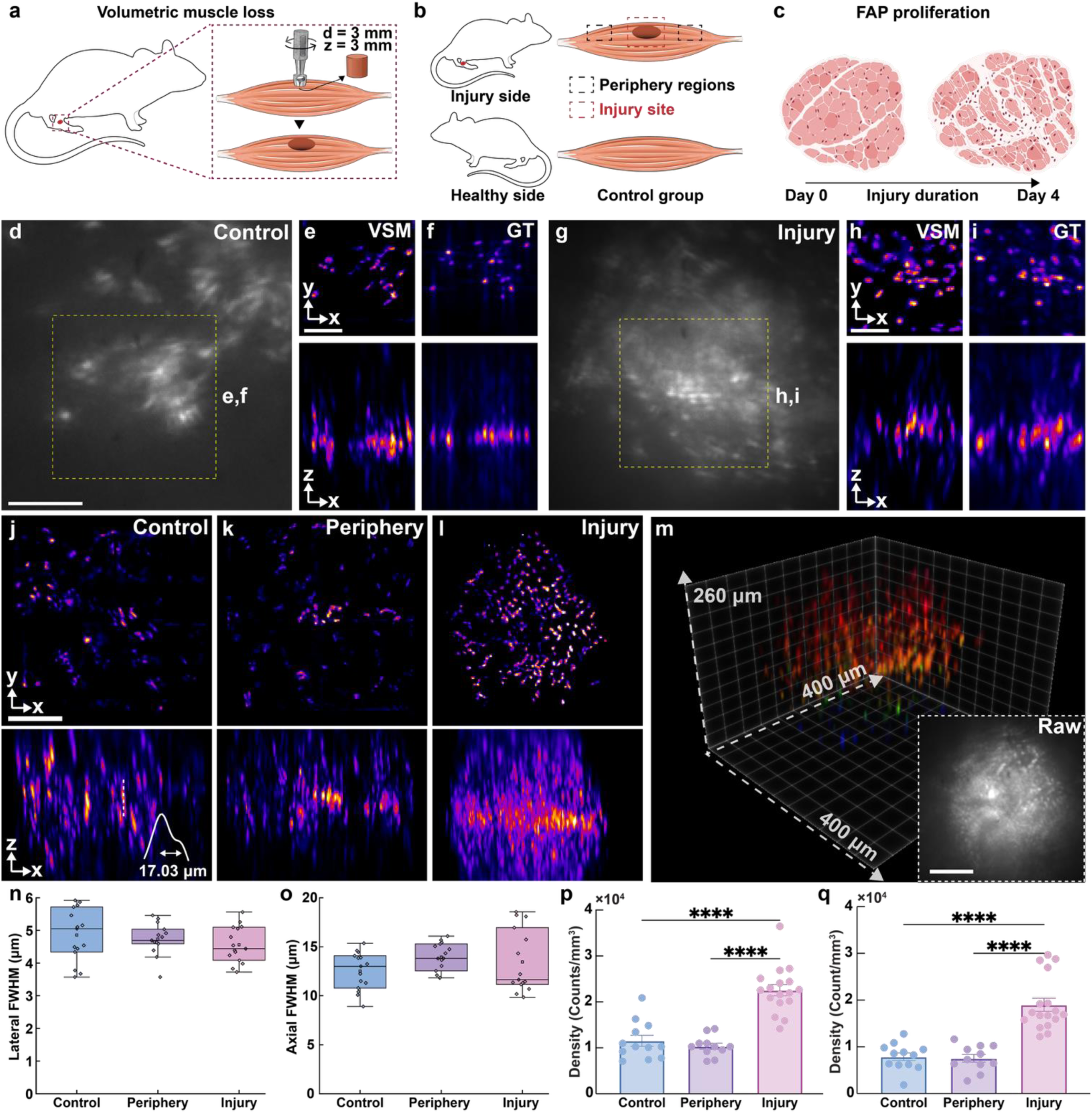
*ex vivo* VSM Imaging of volumetric muscle loss (VML) in the mouse. **(a)** Schematic of the VML model. A cylindrical biopsy (∼3 mm diameter, ∼3 mm depth) was created in the quadriceps muscle to induce the VML. **(b)** Illustration of regions of interest, including the injury site, the peripheral regions located 5-10 mm from the defect, and the contralateral healthy quadriceps serving as a control. A 150-µm-thick mouse skin layer was positioned ∼1 mm above the control and peripheral tissues to introduce layered scattering; at the injury site, the wound depth defines the object-scattering separation. **(c)** Rapid proliferation of FAP after injury, providing essential support for tissue regeneration. **(d)** Representative raw elemental image of the control group, showing sparse FAP nuclei. **(e, f)** Corresponding VSM reconstruction (e), consistent with the ground-truth results (f). **(g)** Representative raw elemental image of the injury group, exhibiting markedly increased FAP density and reduced contrast due to strong scattering. **(h, i)** Corresponding VSM reconstruction (h), showing robust recovery of the three-dimensional nuclear distribution, consistent with the ground-truth results under dense and heterogeneous conditions (f). **(j-l)** VSM images of the control (j), periphery (k), and injury (l) groups with an extended field of view. Volumes were tiled using 3 × 3 stitching with 50% overlap and correlation-based alignment. The intensity profile along the dashed line in (j) resolves two axially nearby nuclei 10-20 µm apart, demonstrating reliable axial discrimination over extended fields. **(m)** Three-dimensional view of the VSM reconstructed injury-site volume, revealing the spatial organization of nuclei with substantially improved clarity relative to the raw measurement (inset). **(n, o)** Lateral (n) and axial (o) resolution analysis across the control, periphery, and injury groups, exhibiting consistent values ranging from 3-6 µm and 12-14 µm, respectively, and the robustness of resolution restoration by VSM from sparse to high-density tissue conditions. **(p, q)** Quantitative FAP density derived from VSM reconstructions (p) and corresponding ground-truth measurements (q). The injury site exhibits an approximately twofold increase in cell density, consistent with injury-induced proliferation. Statistical analyses were performed using ordinary one-way ANOVA followed by Tukey’s multiple comparisons test. ****p < 0.0001. Scale bars: 100 µm.

### *ex vivo* VSM Imaging of Volumetric Muscle Loss in Mouse

Volumetric muscle loss is a severe traumatic condition characterized by the irreversible loss of substantial muscle tissue^60,61^. The substantial defect volume, heterogeneous regenerative microenvironment, and densely remodeled extracellular matrix create a highly scattering optical milieu that significantly hinders high-resolution volumetric imaging. To evaluate VSM under these demanding conditions, we performed *ex vivo* volumetric imaging of fibro-adipogenic progenitors (FAPs), a key stromal population that regulates muscle regeneration^62,63^. Specifically, in the mouse quadriceps volumetric muscle loss model, the defect spans approximately 3 mm laterally and up to 3 mm in depth (**Fig. 3a, Supplementary Fig. 7, Methods)**, forming a structurally complex and optically heterogeneous tissue volume. FAP nuclei were imaged four days post-injury at the injured site and adjacent peripheral regions, with the contralateral healthy quadriceps serving as a control (**Fig. 3b, Supplementary Fig. 7**). As expected from prior studies^64^, FAP proliferation was elevated at the injured site relative to peripheral and control regions (**Fig. 3c)**.

In the control tissue, raw scattering measurements exhibit sparse nuclear distributions (**Fig. 3d**), whereas VSM reconstruction closely matches the corresponding ground truth (**Fig. 3e, f**), demonstrating accurate volumetric recovery under moderate scattering. In contrast, the injured region displays markedly increased cell density, producing denser and lower-contrast speckle patterns in the raw images (**Fig. 3g**). Despite this elevated scattering condition, VSM effectively reconstructs the three-dimensional distribution of FAP nuclei (**Fig. 3h**), achieving strong structural agreement with ground truth measurements (**Fig. 3i**). Furthermore, to assess scalability, we extended VSM to more extensive volumetric coverage by continuous acquisition and stitching multiple reconstructions. Unlike speckle autocorrelation approaches, which degrade when contributions arise outside the OME range, VSM maintains stable reconstruction across tiled fields (**Fig. 3j-l**). For instance, volumetric reconstruction spanning approximately 500 µm laterally resolves axially adjacent nuclei separated by 17 µm (**Fig. 3k**), underscoring the preservation of depth discrimination under high-density conditions. The three-dimensional rendering shown in **Fig. 3m** reinforces the robust performance of VSM, exhibiting high-fidelity reconstruction over a volume of approximately 500 × 500 × 260 µm³. Quantitative analysis further confirms consistent performance across the control, peripheral, and VML tissues, with lateral resolution of approximately 4-5 µm and axial resolution of 12-13 µm (**Fig. 3n, o**). Cell densities derived from VSM reconstructions were 1.15 × 10^4^, 1.03 × 10^4^, and 2.26 × 10^4^ cells/mm^3^ for the control, peripheral, and VML regions, respectively (**Fig. 3p**), closely matching the corresponding ground-truth measurements (**Fig. 3q**). Collectively, these results demonstrate that VSM enables accurate, large-volume three-dimensional reconstruction and preserves quantitative fidelity within highly scattering, biologically complex tissue environments.

### Whole-Organism Volumetric Imaging in *Xenopus* Embryos

Imaging intact whole organisms represents a particularly stringent test for scattering microscopy. Developing embryos contain complex, multilayered tissues with intrinsic pigmentation and strong internal scattering, creating a highly heterogeneous optical environment. Achieving high-resolution, volumetric fluorescence imaging under these conditions without tissue clearing remains a significant challenge.

To advance VSM in this regime, we imaged intact *Xenopus* embryos at three developmental stages (**Fig. 4a**). In this context, fluorescent structures are intrinsically embedded within scattering tissue, requiring robust recovery of volumetric information from strongly distorted measurements. We therefore implemented an extended VSM processing strategy tailored to fully embedded scattering conditions while maintaining the same optical configuration (**Supplementary Fig. 8, Methods**). We first imaged nuclei at the gastrula stage, during which the embryo exhibits an approximately spherical morphology (**Fig. 4b, Methods**). This improvement enabled complete three-dimensional visualization of individual cells throughout the organism (**Fig. 4c**). With volumetric reconstruction, adjacent nuclei separated by ∼5 µm and ∼9 µm were clearly resolved, demonstrating a substantial improvement over the raw light-field reconstruction (**Fig. 4d-g).** The enhanced contrast and effective suppression of scattering background further facilitated markedly improved cell segmentation, in which, for example, we observed 574 nuclei identified with VSM, a more than three-fold improvement compared with 169 objects detected in the unprocessed data, which were severely compromised by scattering artifacts (**Fig. 4h, i**).

**Figure 4.**
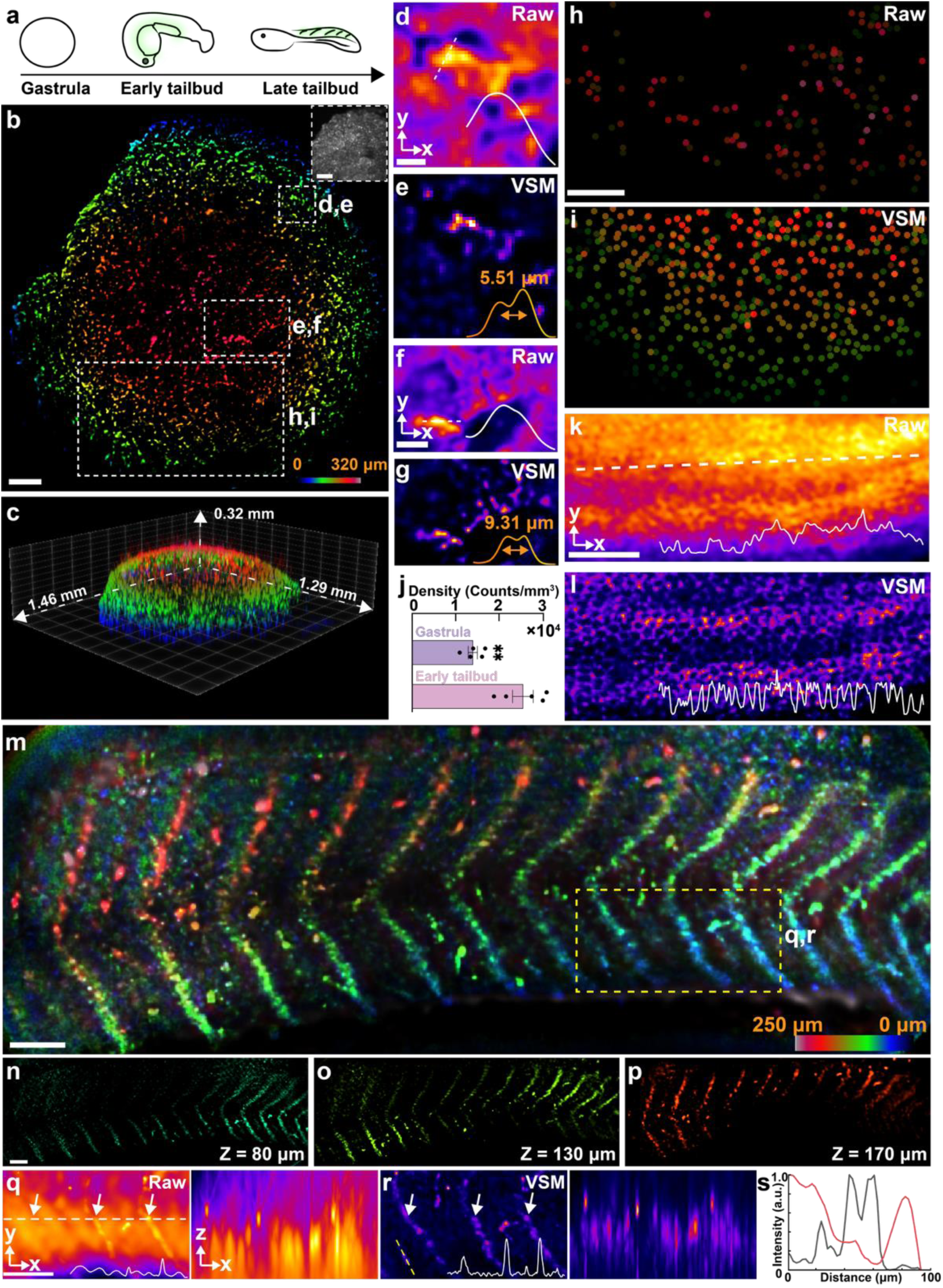
Whole-organism volumetric imaging of *Xenopus* embryos using extended VSM. **(a)** Schematic of *Xenopus* embryo morphology at representative developmental stages. **(b, c)** Depth-color-coded (b) and corresponding three-dimensional perspective (c) images of nuclear distribution at the gastrula stage using VSM, demonstrating high-fidelity nuclear organization with suppressed background scattering over raw elemental images (inset). **(d-g)** Comparative conventional light-field reconstructions (d, f) and VSM-reconstructed volumes (e, g) of regions as marked in (b), resolving adjacent nuclei separated by 5-10 µm with VSM. **(h, i)** Three-dimensional segmentation of nuclei in the boxed region in (b) based on conventional light-field (h) and VSM (i) reconstructions, showing substantially enhanced identification from 169 to 574 nuclei, respectively. **(j)** Nuclear density at the gastrula and early tailbud stages, showing increased cell density during developmental progression. **(k, l)** Conventional light-field reconstructed (k) and VSM (l) images of a caudal spinal cord region at the late tailbud stage, demonstrating the resolution of the epithelial mesh architecture with VSM. **(m-p)** VSM image (m) and individual depth layers (n-p) of the notochord region at the late tailbud stage with an extended field of view at 1.5 × 0.5 × 0.25 mm³, capturing depth-encoded somite morphology. **(q, r)** Comparative raw light-field (q) and VSM (r) reconstructions in *x-y* and *x-z* of the boxed region in (m), revealing adjacent somite architectures in all three dimensions. **(s)** Corresponding cross-sectional intensity profiles along the dashed line in (r), showing recovery of biological features in the VSM reconstruction (black) that are obscured by scattering in the light-field measurement (red). Scale bars: 100 µm (b, h, k, m, n, q), 25 µm (d), 50 μm (f).

We next imaged embryos at the early tailbud stage and quantified nuclear density, observing an approximately two-fold increase relative to the gastrula stage (**Fig. 4j, Supplementary Fig. 9, Methods**), consistent with expected cell proliferation during developmental progression^65^. To further assess performance on continuous anatomical structures, we imaged late tailbud embryos. In the caudal spinal cord, raw light-field reconstruction fails to resolve the mesh-like morphology formed by epithelial cells; the resulting intensity profile lacks distinct, high-contrast features (**Fig. 4k**). Conversely, VSM reconstruction recovers this structure with clarity, revealing well-resolved periodic features (**Fig. 4l**). Notably, by extending VSM to produce millimeter-scale volumes, we generated a 1.5 × 0.5 × 0.25 mm^3^ field of view encompassing the entire notochord region (**Fig. 4m**), where the characteristic V-shaped outline of the somites **(Methods)** is distinctly resolved across multiple axial layers (**Fig. 4n-p**). The results exhibited a restored micrometer-level resolution and approximately two orders of magnitude enhancement in feature recognition with VSM, demonstrating its robust panoramic view of complex, highly scattering organismal tissues (**Fig. 4q-s**).

## DISCUSSION

Biological tissues inherently scatter light, disrupting spatial and angular information and thereby limiting volumetric visualization^66,67^. Despite continued advances, practical three-dimensional imaging through tissue remains constrained by system complexity, restricted volumetric coverage, or axial degeneracy. Accordingly, simple and scan-free strategies for robust volumetric recovery in tissue-relevant environments remain critically needed. In this work, we present volumetric scattering microscopy (VSM), an integrated optical-computational framework for accessible, scan-free three-dimensional imaging in scattering biological media. VSM harnesses the angular information embedded in speckle-encoded measurements to recover volumetric structure without wavefront measurement or mechanical scanning, while preserving the simplicity of a standard epi-fluorescence architecture. The framework enables robust and scalable performance with quantitative accuracy across cellular, tissue, and whole-organism systems, in both layered and fully embedded scattering configurations. Furthermore, the VSM platform is readily extensible through integration with advanced optical designs^68–71^, computational strategies^72–77^, functional imaging probes^78–80^, and versatile instrumental architectures^81–84^. VSM establishes a scalable pathway toward volumetric microscopy in complex biological environments and opens new opportunities for quantitative investigation of dynamic, multiscale biological systems beyond conventional scattering limits.

## METHODS

### Image acquisition

All image acquisitions were performed on a home-built upright Fourier light-field microscopy system equipped with two laser lines (488 nm and 594 nm, Coherent) and a single objective lens (CFI Plan Fluor 20×, 0.5 NA, Nikon). Random speckle illumination was generated using a high-transmittance diffuser (ED1-C20-MD, Thorlabs) mounted on a motorized rotation stage (K10CR2, Thorlabs). The resulting fluorescence signals were collected by the objective lens and relayed by a tube lens (TTL-200A, Thorlabs) to form the native image plane (NIP). The NIP was optically Fourier-transformed using a Fourier lens (ACT508-200-A, Thorlabs), and a customized microlens array (MLA, *f*_MLA_ = 30mm) was positioned at the back focal plane of the Fourier lens. The elemental images formed by the MLA were recorded using an sCMOS camera (Dhyana 400 BSI V3, Tucsen). The resulting elemental images covered a field of view of 600 × 600 µm² with an overall magnification of 3×. During data acquisition, the object is positioned at the focal plane of the objective lens, and a scattering medium is placed above the object at a prescribed distance. After introducing the scattering layer, the focal plane of the objective lens is slightly adjusted to ensure the speckle patterns are centered within each elemental image.

### Robust non-negative principal matrix factorization

The robust non-negative principal matrix factorization (RNP) approach is implemented as a three-stage process based on speckle imagery acquired under longitudinal speckle illumination; in this work, RNP is applied at the elemental-image level. Briefly, the RNP algorithm first preprocesses the image to reduce noise, then separates sparse features from a low-rank, redundant background. Subsequently, the extracted sparse features are reduced in dimension using non-negative matrix factorization. This step enables assigning each speckle-pattern feature to its corresponding emitter, after which the final reconstructed image is obtained by selecting one emitter as a reference and assembling all others based on their relative positional offsets via pairwise Wiener deconvolution. The details of the algorithm are provided in **Supplementary Notes 2 and 3**, and the most up-to-date implementation of RNP is available at https://github.com/ShuJiaLab/VSM.

### Modified VSM framework for imaging objects embedded in an inherently scattering medium

Image acquisition under this condition follows the same procedure as in the deep-scattering environment, where speckle illumination is used to generate dynamic illumination patterns. The captured raw measurements undergo the same pre-processing steps: Gaussian filtering followed by RPCA for feature extraction. In this case, where structural information is weakly discernible and strongly obscured by background scattering, RPCA isolates illumination patterns with minimal background contribution as features. Covariance analysis between the recovered illumination patterns and their corresponding images amplifies the target signal, yielding high-contrast, de-scattered reconstructions^48,85^. The resulting descattered elemental images are then used for volumetric reconstruction using the same sub-pupil alignment and patch-wise deconvolution strategy.

### Preparation of volumetric 2-µm microspheres

An agarose solution was prepared by dissolving 10 mg agarose powder (A9414-10G, Sigma-Aldrich) in 1 mL deionized water (DW) in a plastic tube. 2-µm microspheres were then added to the agarose solution and gently mixed to achieve the desired dilution. The mixture was heated to 80 °C until the solution became fully transparent, indicating complete agarose dissolution. The solution was subsequently vortexed to ensure uniform distribution of the microspheres. Finally, the mixture was dispensed onto a glass slide and stored at 4 °C to allow hydrogel formation prior to imaging.

### Cell membrane staining

Bone marrow–derived MSCs were cultured in the alginate–Ca²⁺ hydrogel droplets. Prior to staining, the culture medium was removed, and the 3D-cultured MSCs were washed three times with PBS. WGA 594 (W11262, Invitrogen) was prepared in PBS at a final concentration of 5 µg/mL. The staining solution was prewarmed to 37 °C and then added to the imaging dish. Cells were incubated with the staining solution for 30 min at 37 °C in a 5% CO₂ atmosphere. After staining, the solution was removed, and the cells were washed three times with PBS. The samples were finally stored in PBS prior to imaging.

### Cell nuclei staining

HeLa cells were cultured in alginate–Ca²⁺ hydrogel droplets and treated with staurosporine (STS) for 0, 4, and 8 h. At each time point, the STS-containing medium was removed, and the cells were washed three times with PBS. SYTO 64 (S11346, Invitrogen) was prepared in PBS at a final concentration of 50 nM. The staining solution was prewarmed to 37 °C and added to the wells. Cells were incubated with the staining solution for 45 min at 37 °C in a 5% CO₂ atmosphere. After staining, the solution was removed, and the cells were washed three times with PBS. The samples were finally stored in PBS prior to imaging.

### Volumetric muscle loss-injured quadriceps tissue preparation

A 24-month-old male B6.129S4-Pdgfratm11(EGFP)Sor/J (PDGFRαEGFP) mouse, in which PDGFRα-expressing cells exhibit enhanced green fluorescent protein (EGFP)-labeled nuclei, was purchased from The Jackson Laboratory and used as a fibro-adipogenic progenitor (FAP) reporter mouse. The volumetric muscle loss (VML) surgical procedure was performed as previously described^86^. Briefly, the mouse was anesthetized with 2% isoflurane, and the left hindlimb was prepped and sterilized. A single skin incision was made over the quadriceps, and a 3 mm biopsy punch was used to create a full-thickness muscle defect at the center of the quadriceps muscle. The skin was closed, and the muscle was allowed to recover without intervention. The mouse was euthanized by CO_2_ inhalation at day 4 post-injury. Both the injured and contralateral quadriceps muscles were harvested and immediately imaged. During imaging, a 150-μm-thick mouse skin was used as the scattering medium. Specifically, for imaging the VML site, the skin was directly placed on top of the muscle without additional spacing, with the intrinsic depth of the injury site serving as the natural separation between the tissue and the scattering layer. For the peripheral and control groups, the same skin sample was positioned approximately 1 mm above the tissue to provide a defined object–scatterer distance.

### *Xenopus* embryo nuclei staining

*Xenopus* embryos at the gastrula and early tailbud stages were used for nuclear staining. Briefly, embryos were fixed in MEMFA for 2 hours at desired stages, rinsed in PBS, and blocked in 10% goat serum diluted in PBT (PBS with 0.1% Triton X-100). The staining solution was prepared by diluting SYTO 64 (S11346, Invitrogen) in PBS to a final concentration of 100 nM, and the embryos were stained for 60 min. After staining, the solution was removed, and the embryos were washed three times with PBS for 5 min each. The samples were then fixed with 4% paraformaldehyde (PFA) for 10 min, followed by three additional PBS washes (5 min each). The samples were finally stored in PBS prior to imaging.

### *Xenopus* tissue staining

*Xenopus* embryos at the late tailbud stage were used for tissue staining. The embryos were fixed, rinsed, and blocked following the same protocol described for nuclei staining. The samples were then incubated with an anti-VE-cadherin antibody (ThermoFisher Scientific, 36-1900) at a 1:50 dilution, followed by a FITC-conjugated secondary antibody at a 1:1000 dilution. Although this staining protocol was intended to label blood vessels, the fluorescence signal was predominantly observed in the somite regions, forming characteristic V-shaped patterns. In addition, mesh-like fluorescence structures were detected in the tail region, suggesting association with epithelial structures.

### Chemical and biological materials

The sources of the chemicals and biological materials used in the experiments, including company names and catalog numbers, are listed in **Supplementary Table 3**. No further authentication procedures are conducted after receiving the stock cell lines directly from the suppliers.

## DATA AVAILABILITY

The data sets generated and analyzed in this study are available from the corresponding author upon request.

## CODE AVAILABILITY

The code has been written in MATLAB (MathWorks) and tested with versions 2022a and 2022b. The most updated version of the software can be found at https://github.com/ShuJiaLab/VSM upon publication.

## ACKNOWLEDGEMENTS

We acknowledge support from National Science Foundation grants 2503686, 2225990, and 2145235, National Institutes of Health grants R35GM124846 and R21HD110918, and the faculty start-up fund of the Georgia Institute of Technology.

## AUTHOR CONTRIBUTIONS STATEMENT

Z.G. and S.J. conceived and designed the project. Z.G., Z.L., and K.H. contributed to the construction of the optical system. Z.G. and K.H. conducted image processing and developed algorithms. Z.G., H.Z., E.A.B and S.N. prepared biological samples. Z.G. performed imaging experiments. S.J. supervised the overall project. Z.G. and S.J. wrote the manuscript with input from all authors.

## REFERENCES

1 Schermelleh, L. et al. Super-resolution microscopy demystified. Nature cell biology 21, 72–84 (2019).

2 Yun, S. H. & Kwok, S. J. Light in diagnosis, therapy and surgery. Nature biomedical engineering 1, 0008 (2017).

3 Wu, J., Ji, N. & Tsia, K. K. Speed scaling in multiphoton fluorescence microscopy. Nature Photonics 15, 800–812 (2021).

4 Mikami, H., Gao, L. & Goda, K. Ultrafast optical imaging technology: principles and applications of emerging methods. Nanophotonics 5, 497–509 (2016).

5 Bond, C., Santiago-Ruiz, A. N., Tang, Q. & Lakadamyali, M. Technological advances in super-resolution microscopy to study cellular processes. Molecular Cell 82, 315–332 (2022).

6 Prakash, K. et al. Resolution in super-resolution microscopy—definition, trade-offs and perspectives. Nature Reviews Molecular Cell Biology 25, 677–682 (2024).

7 Liu, S., Hoess, P. & Ries, J. Super-resolution microscopy for structural cell biology. Annual review of biophysics 51, 301–326 (2022).

8 Li, S. et al. AI-empowered super-resolution microscopy: a revolution in nanoscale cellular imaging. Nature Methods, 1–23 (2025).

9 Steelman, Z. A., Ho, D. S., Chu, K. K. & Wax, A. Light-scattering methods for tissue diagnosis. Optica 6, 479–489 (2019).

10 Tuchin, V. V. (Society of Photo-Optical Instrumentation Engineers (SPIE) Bellingham, WA, USA).

11 Yoon, S. et al. Deep optical imaging within complex scattering media. Nature Reviews Physics 2, 141–158 (2020).

12 Bertolotti, J. & Katz, O. Imaging in complex media. Nature Physics 18, 1008–1017 (2022).

13 Cao, H., Mosk, A. P. & Rotter, S. Shaping the propagation of light in complex media. Nature Physics 18, 994–1007 (2022). 10.1038/s41567-022-01677-x

14 Xu, C., Nedergaard, M., Fowell, D. J., Friedl, P. & Ji, N. Multiphoton fluorescence microscopy for in vivo imaging. Cell 187, 4458–4487 (2024).

15 Ou, Z. et al. Achieving optical transparency in live animals with absorbing molecules. Science 385, eadm6869 (2024).

16 Yu, T., Zhu, J., Li, D. & Zhu, D. Physical and chemical mechanisms of tissue optical clearing. Iscience 24 (2021).

17 Ueda, H. R. et al. Tissue clearing and its applications in neuroscience. Nature Reviews Neuroscience 21, 61–79 (2020).

18 Prevedel, R. et al. Three-photon microscopy: an emerging technique for deep intravital brain imaging. Nature Reviews Neuroscience 26, 521–537 (2025).

19 Papagiakoumou, E., Ronzitti, E. & Emiliani, V. Scanless two-photon excitation with temporal focusing. Nature Methods 17, 571–581 (2020).

20 Maruca, S., Rehain, P., Sua, Y. M., Zhu, S. & Huang, Y. Non-invasive single photon imaging through strongly scattering media. Optics Express 29, 9981–9990 (2021).

21 Kang, P. et al. Optical transfer function of time-gated coherent imaging in the presence of a scattering medium. Optics Express 29, 3395–3405 (2021).

22 Jeong, S. et al. Focusing of light energy inside a scattering medium by controlling the time-gated multiple light scattering. Nature Photonics 12, 277–283 (2018).

23 Ko, H. et al. Acousto-optic volumetric gating for reflection-mode deep optical imaging within a scattering medium. ACS Photonics 10, 3664–3673 (2023).

24 Jang, M. et al. Deep tissue space-gated microscopy via acousto-optic interaction. Nature communications 11, 710 (2020).

25 Ruan, H., Liu, Y., Xu, J., Huang, Y. & Yang, C. Fluorescence imaging through dynamic scattering media with speckle-encoded ultrasound-modulated light correlation. Nature Photonics 14, 511–516 (2020).

26 Katz, O., Ramaz, F., Gigan, S. & Fink, M. Controlling light in complex media beyond the acoustic diffraction-limit using the acousto-optic transmission matrix. Nature communications 10, 717 (2019).

27 Yu, Z., et al. Wavefront shaping: a versatile tool to conquer multiple scattering in multidisciplinary fields. The Innovation 3 (2022).

28 Rotter, S. & Gigan, S. Light fields in complex media: Mesoscopic scattering meets wave control. Reviews of Modern Physics 89, 015005 (2017).

29 Gigan, S. et al. Roadmap on wavefront shaping and deep imaging in complex media. Journal of Physics: Photonics 4, 042501 (2022).

30 Kim, M., Choi, W., Choi, Y., Yoon, C. & Choi, W. Transmission matrix of a scattering medium and its applications in biophotonics. Optics express 23, 12648–12668 (2015).

31 Boniface, A., Dong, J. & Gigan, S. Non-invasive focusing and imaging in scattering media with a fluorescence-based transmission matrix. Nature communications 11, 6154 (2020).

32 Baek, Y., de Aguiar, H. B. & Gigan, S. Three-dimensional holographic imaging of incoherent objects through scattering media. Nature Communications (2025).

33 Lindell, D. B. & Wetzstein, G. Three-dimensional imaging through scattering media based on confocal diffuse tomography. Nature communications 11, 4517 (2020).

34 Gigan, S. Imaging and computing with disorder. Nature Physics 18, 980–985 (2022).

35 Bertolotti, J. et al. Non-invasive imaging through opaque scattering layers. Nature 491, 232– 234 (2012).

36 Katz, O., Heidmann, P., Fink, M. & Gigan, S. Non-invasive single-shot imaging through scattering layers and around corners via speckle correlations. Nature photonics 8, 784–790 (2014).

37 Freund, I., Rosenbluh, M. & Feng, S. Memory effects in propagation of optical waves through disordered media. Physical review letters 61, 2328 (1988).

38 Hofer, M., Soeller, C., Brasselet, S. & Bertolotti, J. Wide field fluorescence epi-microscopy behind a scattering medium enabled by speckle correlations. Optics express 26, 9866–9881 (2018).

39 Osnabrugge, G., Horstmeyer, R., Papadopoulos, I. N., Judkewitz, B. & Vellekoop, I. M. Generalized optical memory effect. Optica 4, 886–892 (2017).

40 Kang, S. et al. High-resolution adaptive optical imaging within thick scattering media using closed-loop accumulation of single scattering. Nature Communications 8, 2157 (2017). 10.1038/s41467-017-02117-8

41 Weinberg, G., Sunray, E. & Katz, O. Noninvasive megapixel fluorescence microscopy through scattering layers by a virtual incoherent reflection matrix. Sci Adv 10, eadl5218 (2024). 10.1126/sciadv.adl5218

42 Li, Y., Xue, Y. & Tian, L. Deep speckle correlation: a deep learning approach toward scalable imaging through scattering media. Optica 5, 1181–1190 (2018).

43 Liu, H. et al. Learning-based real-time imaging through dynamic scattering media. Light: Science & Applications 13, 194 (2024). 10.1038/s41377-024-01569-0

44 Alido, J. et al. Robust single-shot 3D fluorescence imaging in scattering media with a simulator-trained neural network. Optics Express 32, 6241–6257 (2024).

45 Xu, X. et al. Descattering and image restoration with a transformer-based neural network in deep tissue imaging. Proceedings of the National Academy of Sciences 122, e2503576122 (2025).

46 Premillieu, E., Caravaca-Aguirre, A. M., Labouesse, S., Irsch, K. & Piestun, R. Aberration correction in epi-fluorescence microscopy using unknown speckle illumination. Optics Express 32, 37658–37667 (2024).

47. Mangeat, T., et al. Super-resolved live-cell imaging using random illumination microscopy. Cell Reports Methods 1 (2021).

48 Taylor, M. A., Nöbauer, T., Pernia-Andrade, A., Schlumm, F. & Vaziri, A. Brain-wide 3D light-field imaging of neuronal activity with speckle-enhanced resolution. Optica 5, 345–353 (2018).

49 Moretti, C. & Gigan, S. Readout of fluorescence functional signals through highly scattering tissue. Nature Photonics 14, 361–364 (2020). 10.1038/s41566-020-0612-2

50 Zhu, L. et al. Large field-of-view non-invasive imaging through scattering layers using fluctuating random illumination. Nat Commun 13, 1447 (2022). 10.1038/s41467-022-29166-y

51 Gao, Z. et al. Fluorescence microscopy through scattering media with robust matrix factorization. Cell Rep Methods 5, 101031 (2025). 10.1016/j.crmeth.2025.101031

52 Okamoto, Y., Horisaki, R. & Tanida, J. Noninvasive three-dimensional imaging through scattering media by three-dimensional speckle correlation. Optics letters 44, 2526–2529 (2019).

53 Aarav, S. & Fleischer, J. W. Depth-resolved speckle correlation imaging using the axial memory effect. Optics Express 32, 23750–23757 (2024).

54 Horisaki, R., Okamoto, Y. & Tanida, J. Single-shot noninvasive three-dimensional imaging through scattering media. Optics letters 44, 4032–4035 (2019).

55 Goodman, J. W. Speckle phenomena in optics: theory and applications. (Roberts and Company Publishers, 2007).

56 Ling, Z. et al. Multiscale and recursive unmixing of spatiotemporal rhythms for live-cell and intravital cardiac microscopy. Nat Cardiovasc Res 4, 637–648 (2025). 10.1038/s44161-025-00649-7

57 Liu, W., Kim, G.-A. R., Takayama, S. & Jia, S. Fourier light-field imaging of human organoids with a hybrid point-spr ead function. Biosensors and Bioelectronics 208, 114201 (2022).

58. Gao, Z., et al. Fluorescence microscopy through scattering media with robust matrix factorization. Cell Reports Methods 5 (2025).

59 Fu, B. et al. Patch deconvolution for Fourier light-field microscopy. bioRxiv, 2025.2006. 2013.659385 (2025).

60 Grogan, B. F., Hsu, J. R. & Consortium, S. T. R. Volumetric muscle loss. JAAOS-Journal of the American Academy of Orthopaedic Surgeons 19, S35–S37 (2011).

61 Hu, C. et al. A mouse model of volumetric muscle loss and therapeutic scaffold implantation. Nature protocols 20, 608–619 (2025).

62 Joe, A. W. et al. Muscle injury activates resident fibro/adipogenic progenitors that facilitate myogenesis. Nature cell biology 12, 153–163 (2010).

63 Loomis, T. & Smith, L. R. Thrown for a loop: fibro-adipogenic progenitors in skeletal muscle fibrosis. American Journal of Physiology-Cell Physiology 325, C895–C906 (2023).

64 Anderson, S. E. et al. Aberrant Fibro-Adipogenic Progenitor Subpopulations Drive Volumetric Muscle Loss-Induced Fibrosis. bioRxiv, 2025.2005. 2011.653339 (2025).

65 Cooke, J. Cell number in relation to primary pattern formation in the embryo of Xenopus laevis: I. The cell cycle during new pattern formation in response to implanted organizers. Development 51, 165–182 (1979).

66 Jacques, S. L. Optical properties of biological tissues: a review. Physics in Medicine & Biology 58, R37 (2013).

67 Andreou, C., Weissleder, R. & Kircher, M. F. Multiplexed imaging in oncology. Nature Biomedical Engineering 6, 527–540 (2022). 10.1038/s41551-022-00891-5

68 Zhu, L. et al. Large field-of-view non-invasive imaging through scattering layers using fluctuating random illumination. Nature communications 13, 1447 (2022).

69 Haim, O., Boger-Lombard, J. & Katz, O. Image-guided computational holographic wavefront shaping. Nature Photonics 19, 44–53 (2025). 10.1038/s41566-024-01544-6

70 Wang, R. et al. Multiscale aperture synthesis imager. Nature Communications 16, 10582 (2025). 10.1038/s41467-025-65661-8

71 Liang, H. et al. An optical meta-image-processor for enhanced imaging through strongly scattering media. Nature Communications 16, 9732 (2025).

72 Tahir, W., Wang, H. & Tian, L. Adaptive 3D descattering with a dynamic synthesis network. Light: Science & Applications 11, 42 (2022).

73 Barbastathis, G., Ozcan, A. & Situ, G. On the use of deep learning for computational imaging. Optica 6, 921–943 (2019).

74 Kwon, T. et al. Video Diffusion Posterior Sampling for Seeing Beyond Dynamic Scattering Layers. IEEE Transactions on Pattern Analysis and Machine Intelligence 47, 11152–11167 (2025). 10.1109/TPAMI.2025.3598457

75 Sunray, E., Weinberg, G., Laufer, B. & Katz, O. Matrix-based imaging through dynamic scattering. Nature Communications 16, 9413 (2025).

76 Han, K. et al. Volumetric localization microscopy with deep learning. Nature Communications 16, 10960 (2025). 10.1038/s41467-025-65941-3

77 Mandracchia, B. et al. Optimal sparsity allows reliable system-aware restoration of fluorescence microscopy images. Sci Adv 9, eadg9245 (2023). 10.1126/sciadv.adg9245

78 Defienne, H. et al. Advances in quantum imaging. Nature Photonics 18, 1024–1036 (2024). 10.1038/s41566-024-01516-w

79 Chen, Y., Wang, S. & Zhang, F. Near-infrared luminescence high-contrast in vivo biomedical imaging. Nature Reviews Bioengineering 1, 60–78 (2023).

80 Liu, K. et al. Deep and dynamic metabolic and structural imaging in living tissues. Science Advances 10, eadp2438 (2024). doi:10.1126/sciadv.adp2438

81 Linda Liu, F., Kuo, G., Antipa, N., Yanny, K. & Waller, L. Fourier DiffuserScope: single-shot 3D Fourier light field microscopy with a diffuser. Optics Express 28, 28969–28986 (2020). 10.1364/OE.400876

82 Wang, Z. et al. Kilohertz volumetric imaging of in vivo dynamics using squeezed light field microscopy. Nature Methods 22, 2194–2204 (2025). 10.1038/s41592-025-02843-8

83 Yang, J. et al. An ultrahigh-fidelity 3D holographic display using scattering to homogenize the angular spectrum. Science Advances 9, eadi9987 (2023). doi:10.1126/sciadv.adi9987

84 Qian, C. et al. Breaking the fundamental scattering limit with gain metasurfaces. Nature Communications 13, 4383 (2022). 10.1038/s41467-022-32067-9

85 Gao, Z., Han, K., Hua, X., Liu, W. & Jia, S. hydro SIM: super-resolution speckle illumination microscopy with a hydrogel diffuser. Biomedical Optics Express 15, 3574–3585 (2024).

86 Hymel, L. A. et al. Identifying dysregulated immune cell subsets following volumetric muscle loss with pseudo-time trajectories. Communications Biology 6, 749 (2023).

